# Plasticity of the mutation rate and spectrum in eukaryotic marine phytoplankton

**DOI:** 10.64898/2026.07.24.739820

**Authors:** Lisa Mettrop, Anna Lipzen, Celine Vandecasteele, Camille Eché, Frédéric Sanchez, Anaïs Labécot, Eric Manirakiza, Kerrie Barry, Igor V. Grigoriev, Gwenaël Piganeau, Marc Krasovec

## Abstract

The ability to adapt shapes the future of any species or population. Adaptive potential is primarily determined by the spontaneous mutation rate *µ* and the mutation spectrum, which define the new genetic variation available to selection. Increasing evidence points to the plasticity of the mutation rate and spectrum, which vary according to environment. However, data is lacking for marine phytoplankton, an essential group for marine ecosystems as well as a major player in biogeochemical cycles. Here, we measure the plasticity of mutation rates and spectra in the chlorophyte *Ostreococcus tauri* by mutation accumulation experiments in 4 different conditions: low salinity, high salinity, low temperature, and high temperature. *µ_SNM_* is lowest under low salinity, doubles under high salinity and low temperature and quadruples under high temperature. SNM spectra become increasingly biased towards C:G→T:A transitions with increasing *µ_SNM_*. While *µ_ID_* does not change with the environment, the structural mutation rate *µ_SM_* is higher at low temperature, likely due to transposon activity. Given the rapid environmental changes associated with global climate change, mutation rates and spectra plasticity may critically influence the adaptive potential of marine primary producers.

## INTRODUCTION

Climate change drives rapid environmental shifts with major consequences for terrestrial and aquatic ecosystems. Aquatic species increasingly face extreme conditions such as heatwaves^1^, salinity extremes^2^, eutrophication^3,4^ and acidification^5^. While most species have some phenotypic plasticity that allows them to survive stresses up to a certain point^6^, adaptation comes from de novo mutations. The spontaneous mutation rate (*µ*), together with the mutation spectrum, define the quantity and type of new genetic material available to selection. Thus, *µ* has long been a focus of evolutionary biology^7–11^. Overall, *µ* is an evolvable trait that follows the Drift Barrier Hypothesis (DBH)^12^: *µ* decreases under the effect of selection, reducing the deleterious mutation load until the fitness gain of further decreasing *µ* will have a selection coefficient too small to be noticeable. At this point, *µ* is defined by genetic drift. In populations with a large effective population size (*N_e_*), selection is more efficient and *µ* is expected to be lower than in small populations.

The standard for *µ* estimations in unicellular species is the mutation accumulation (MA) experiment^13^. It starts with a clonal population from which MA lines are started. At regular intervals, the MA lines are submitted to single-cell bottlenecks, keeping *N_e_*very low and selection negligible; mutations can therefore be fixed by genetic drift independently of their fitness effect. After a large number of generations, the genomes of the MA lines are sequenced and compared to the parent genome to identify de novo mutations. Mutation rate estimates are now available for about 200 species across the tree of life^14,15^.

Furthermore, it has been known for almost a century that *µ* changes with the environment^11,16^, but relatively few studies have been done to document *µ* plasticity in eukaryotes (Table S1). In *A. thaliana*, the single nucleotide mutation rate (*µ_SNM_*) and insertion-deletion rate (*µ_ID_*) go up when temperature or salinity increases^17,18^, while decreasing temperature also changes *µ*, indicating that any distance from standard temperature increases *µ*^19^. Similarly, in mosquitos, mutation rate varies by 5 when temperature moves away from the standard temperature, both in higher and lower temperatures^17,20^. Another example is the effect of growth media in yeast^21,22^, which acts on both mutation rate and spectrum. Mutation spectrum variation with environment, also observed in *C. reinhardtii*^23^, may be a key of adaptation by exploring previously unsampled mutations^24^. Such variations of mutation rate and spectrum suggest that the environment may be a key determinant of mutation rate, on par with *N_e_* or other biological traits, shaping evolutionary trajectories among populations living in different ecological contexts.

In the present work, we evaluate the variation of mutation rate and spectrum with environmental parameters in the green alga *Ostreococcus tauri*, the smallest free- living eukaryote known, with a gene-dense genome of 13Mb for a GC content of 59.5%^25^. This cosmopolitan alga, part of Mamiellophyceae, is a model species for phytoplankton in coastal areas. We performed MA experiments in four different conditions: high salinity (50 PSU), low salinity (20 PSU), high temperature (26°C) and low temperature (14°C).

## RESULTS

### Mutation accumulation experiments

The MA experiments lasted between 210 and 336 days, with 15 to 24 bottlenecks (Table S2). Total accumulated generations were 9,287 for low salt condition (‘S20’), 12,041 for high salt condition (‘S50’), 10,973 for low temperature (‘T14’) and 4,304 for high temperature (‘T26’) (Tables 1, S2 and S3). The difference in total generations is mostly due to the losses of MA lines during the experiments. The callable genome (*G\**) per MA ranged from 9,358,358 bp in T26 (71.8% of the reference genome) to 11,641,979 in T14 (89.3% of the reference). *G\** was not evenly divided over the chromosomes (Table S4), which is mostly due to the particular genome architecture of the species. *O. tauri* features two ‘outlier’ chromosomes: The big outlier chromosome (‘BOC’, Chr2) and the small outlier chromosome (‘SOC’, Chr19)^25^. These chromosomes have the particularity of a lower GC-content than the rest of the genome with more repetitive sequences. In each MA, the BOC and the SOC were less than half covered (on average 48% and 39% coverage, respectively). The organelle genomes were underrepresented in *G\** as well, on average 62% for the chloroplast and 42% for the mitochondrion (Table S4). There was a significant difference of growth rate (GR) in control cultures between the different MA conditions (Fig. 1), with lower GR of 1.77 at S50 (pairwise Wilcoxon rank sum test, p-value < 0.006 for all). T26 grew the fastest, with GR of 2.01 and was significantly faster than T14 (GR = 1.88, Wilcoxon rank sum test, p-value = 0.01) but not than S20 (GR = 1.96, Wilcoxon rank sum test, p-value = 0.64).

**Figure 1.**
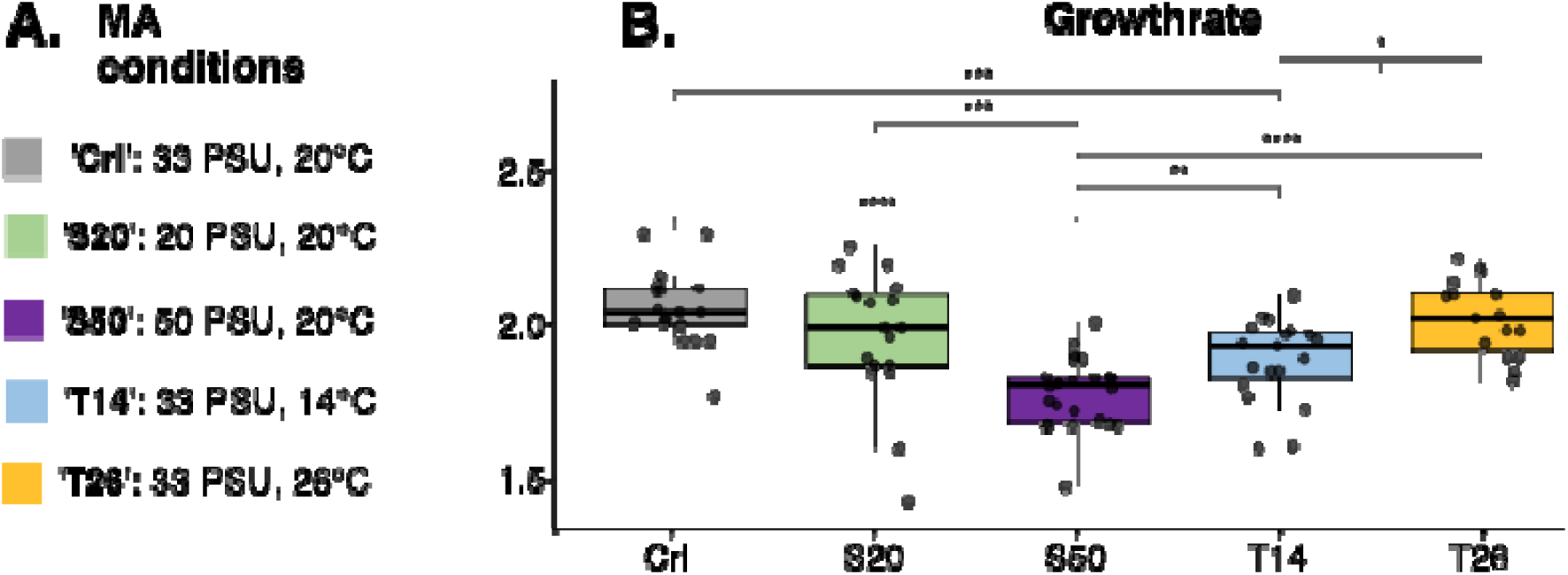
Growth rate (GR) of control cultures in mutation accumulation (MA) experiment conditions. **A**. MA conditions, either salinity or temperature differs from standard lab conditions (‘Crl’). S20: low salinity, S50: high salinity, T14: low temperature, T26: high temperature. **B**. GR of control cultures. Control cultures did not undergo single-cell but 100-cell bottlenecks every two weeks. Each bottleneck the GR was inferred from cell count. GR in high salt conditions (S50) is significantly lower than all other conditions; GR in low temperature (T14) was also lower than in high temperature (T26). Differences were tested with a pairwise Wilcoxon rank sum tests.

### Single nucleotide mutations and small insertions and deletions

A total of 340 nucleotide mutations have been identified: 40 in S20, 123 in S50, 110 in T14 and 67 in T26 (Table S5). Nuclear single nucleotide mutation rates (*µ_SNM_*) differed between MA experiments (Table 1, Fig. 2) from a minimum in S20 (similar to the *µ_SNM_*in standard lab conditions^26^) to a maximum in T26 which is 4 times higher than standard lab conditions. S50 and T14 had a *µ_SNM_* both ±2 times higher than standard lab conditions.

**Figure 2.**
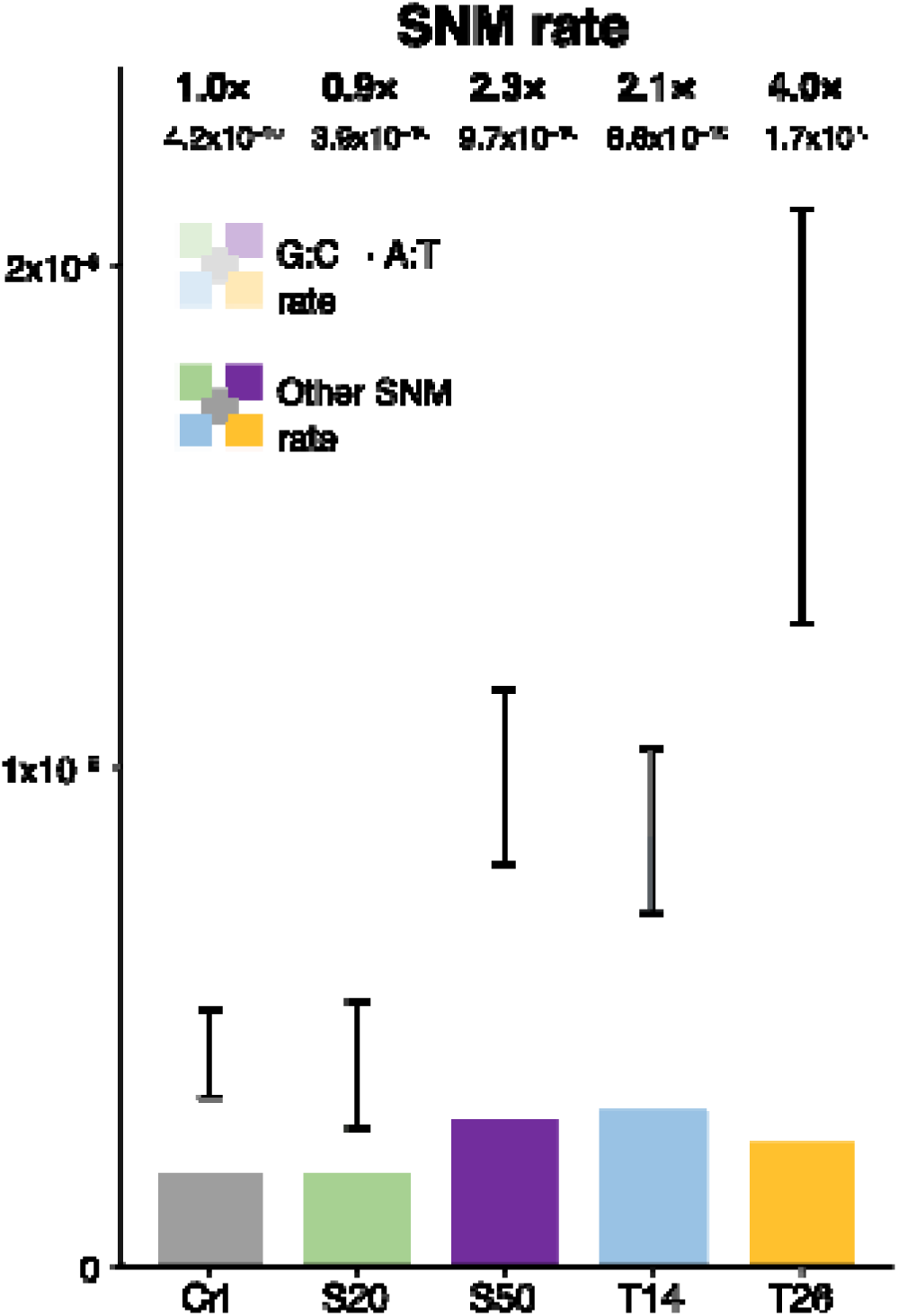
Comparison of *µ_SNM_*between MA conditions shows plasticity. Total bars represent *µ_SNM_* in SNM/basepair/generation and error bars represent the Poisson 95% confidence interval of this total rate. *µ_SNM_* in each condition is marked above the bar, as well as the fold change as compared to standard lab conditions^26^. Thus, *µ_SNM_* remains unchanged in low salt conditions (S20), but doubles in high salt conditions (S50) and low temperature (T14), and quadruples in high temperature (T26). Interestingly, this difference can be attributed for the most part to G:C→A:T mutations, which increase with *µ_SNM_* in a linear fashion (Fig. S2). The proportion of G:C→A:T for each condition is shown in light color in each bar and accounts for 83% of SNM in T26.

**Table 1.** Details of SNM. . Gen.: generations, CDS: coding sequences, Syn:Nsyn: the ratio synonymous to non-synonymous SNM in coding sequences, Ts: transistions, Tv: transversions, (CI95%): Poisson distribution 95% confidence interval, *µ_Ts_*: transitions/*G\**/generation, *µ_Ts_* C:G→T:A: (number of C→T+G→A mutations)/(GC positions in *G\**)/generation. Also see table S2.

| MA | MA lines | Total gen. | SNM | SNM in CDS | Syn: Nsyn | $\mu_{Ts}$ (CI95%) | $\mu_{Ts}$ C:G→T:A (CI95%) | $\mu_{Ts}$ T:A→C:G (CI95%) | $\mu_{Tv}$ (CI95%) | Ts:Tv | Total AT-bias | Transi-tion AT-bias |
| --- | --- | --- | --- | --- | --- | --- | --- | --- | --- | --- | --- | --- |
| S20 | 43 | 9287 | 40 | 35 | 0.52 | 2.53x10<sup>-10</sup><br>(1.66 - 3.71x10<sup>-10</sup>) | 3.40x10<sup>-10</sup><br>(2.10 - 5.20x10<sup>-10</sup>) | 1.22x10<sup>-10</sup><br>(3.98x10<sup>-11</sup> - 2.86x10<sup>-10</sup>) | 1.36x10<sup>-10</sup><br>(7.46x10<sup>-11</sup> - 2.29x10<sup>-10</sup>) | 1.86 | 2.64 | 2.79 |
| S50 | 45 | 12041 | 123 | 92 | 0.61 | 8.20x10<sup>-10</sup><br>(6.70 - 9.93x10<sup>-10</sup>) | 1.13x10<sup>-9</sup><br>(9.03x10<sup>-10</sup> - 1.39x10<sup>-9</sup>) | 3.55x10<sup>-10</sup><br>(2.10 - 5.61x10<sup>-10</sup>) | 1.50x10<sup>-10</sup><br>(9.02x10<sup>-11</sup> - 2.34x10<sup>-10</sup>) | 5.47 | 3.2 | 3.18 |
| T14 | 41 | 10973 | 110 | 95 | 0.4 | 7.28x10<sup>-10</sup><br>(5.88 - 8.92x10<sup>-10</sup>) | 9.17x10<sup>-10</sup><br>(7.15x10<sup>-10</sup> - 1.16E-09) | 4.47x10<sup>-10</sup><br>(2.84 - 6.71x10<sup>-10</sup>) | 1.33x10<sup>-10</sup><br>(7.75x10<sup>-11</sup> - 2.13x10<sup>-10</sup>) | 5.47 | 1.89 | 2.05 |
| T26 | 26 | 4304 | 67 | 55 | 0.53 | 1.49x10<sup>-9</sup><br>(1.13 - 1.92x10<sup>-9</sup>) | 2.34x10<sup>-9</sup><br>(1.77 - 3.03x10<sup>-9</sup>) | 1.89x10<sup>-10</sup><br>(3.89x10<sup>-11</sup> - 5.51x10<sup>-10</sup>) | 1.74x10<sup>-10</sup><br>(6.99x10<sup>-11</sup> - 3.58x10<sup>-10</sup>) | 8.57 | 5.41 | 12.38 |
| Krasov<br>ec et<br>al. | 40 | 17250 | 91 | 61 | 0.49 | 3.08x10<sup>-10</sup><br>(2.39 - 3.92x10<sup>-10</sup>) | / | / | 1.11x10<sup>-10</sup><br>(7.08x10<sup>-11</sup> - 1.64x10<sup>-10</sup>) | 2.79 | 1.74 | / |
| 2017 |  |  |  |  |  |  |  |  |  |  |  |  |

Each MA showed a bias towards transitions (Table 1). The Ts:Tv ratio varied between MA experiments but not significantly (Fisher exact test, p-value > 0.3 for all). Only the rate of C:G→A:T transitions (C→T and G→A) changes between MA experiments and reflects the difference in *µ_SNM_* (Fig. 2, Fig. S1). *µ_C:G→T:A_* is 2.7, 3.3 and 6.9 times higher in T14, S50 and T26 than in S20, respectively (Table 1); in T26, it constitutes 83% of total *µ_SNM_*, while in S50 and T14, *µ_C:G_*_→_*T:A* corresponds to 70% and 64% of total *µ_SNM_*, respectively (Fig. 2). *µ_C:G_*_→_*T:A* in S20 is significantly different from all other samples (Fisher’s exact test, p-value < 10^-5^ for all). We tested the relationship between *µ_C:G_*_→_*T:A* and total *µ_SNM_*across the four MA experimental conditions: Pearson’s correlation showed a strong, significant linear relationship (r = 0.995, p = 0.005), which was supported by linear regression (adjusted R² = 0.985, p = 0.005) (Fig. S2). These results suggest that C:G→T:A mutations explain the majority of the increase in *µ_SNM_* across conditions.

Further, we looked at the AT-bias, defined as the ratio of mutation rates *µ_GC_*_→_*AT*/*µ_AT_*_→_*GC* (Table 1), which impacts genome GC content. The global AT-bias ranged from 1.89 in T14 to 5.41 in T26 (Table 1) but was not significantly different between conditions (Fisher exact test, p-value > 0.4 for all). In addition to the AT- biased mutation rate, the number of mutations from GC to AT and reverse was also not equal (one-sided binomial test of *N_GC_*_→_*AT* on *N_GC_*_→_*AT+AT*_→_*GC*, 60% probability, p-value < 0.02 for all MA conditions).

Last, there was 1 multi-nucleotide mutation in T14 (ACTAGGT→TGTGCGC) (µMN = 7.83×10-12).

We also calculated small insertion-deletion (indel) rates (*µ_ID_*), with 2 events in S20, 8 events in S50, 6 events in T14 and 1 event in T26 (Table S5). Contrary to *µ_SNM_*, there is no significant difference of *µ_ID_*between standard lab and tested MA conditions (Table S2), but the small number of events prevents robust statistical tests.

Regarding the distribution of mutations in the different genomic compartments, *µ* stayed similar between coding sequences (CDS), introns and intergenic regions (Tables S6, S7); only in S50, *µ_SNM_* was significantly higher in intergenic regions than in CDS (Fisher exact test, p-value = 0.002). To test for selection, we compared the synonymous and non-synonymous mutation rates, assuming that one third of positions are synonymous. There was no evidence for a different mutation rate between non-synonymous and synonymous sites (Binomial test, N_syn_mutations_:N_SNM_CDS_, p-value > 0.33 for all) and thus no evidence for selection in any MA, as expected given the low *N_e_* maintained during the experiment.

Finally, we looked at mutations in organelles, with only one multi-nucleotide mutation that seems to be a small inversion in the chloroplast in T26 (AAAG→CTTT). This event was confirmed by PCR amplification and Sanger sequencing of the mutated region. With 1 event, the estimated *µ_chloro_* at 26°C is 4.31×10^-9^ (CI95% 1.09×10^-10^- 2.40×10^-8^) (Table S2).

### Whole chromosome duplications (WCD)

A total 12 WCD in 11 different MA lines were observed (Fig. 3 and S3, Table S8). Chr4, Chr11 and Chr13 were duplicated twice in different MA lines, while many other chromosomes were never duplicated. WCD rates per genome per generation (*U_WCD_*) was the lowest in S50, followed by increasing rates in S20, T14 and T26. However, the difference in *U_WCD_* was not significant (One-sided Fisher exact test, p-value > 0.05 for all).

**Figure 3.**
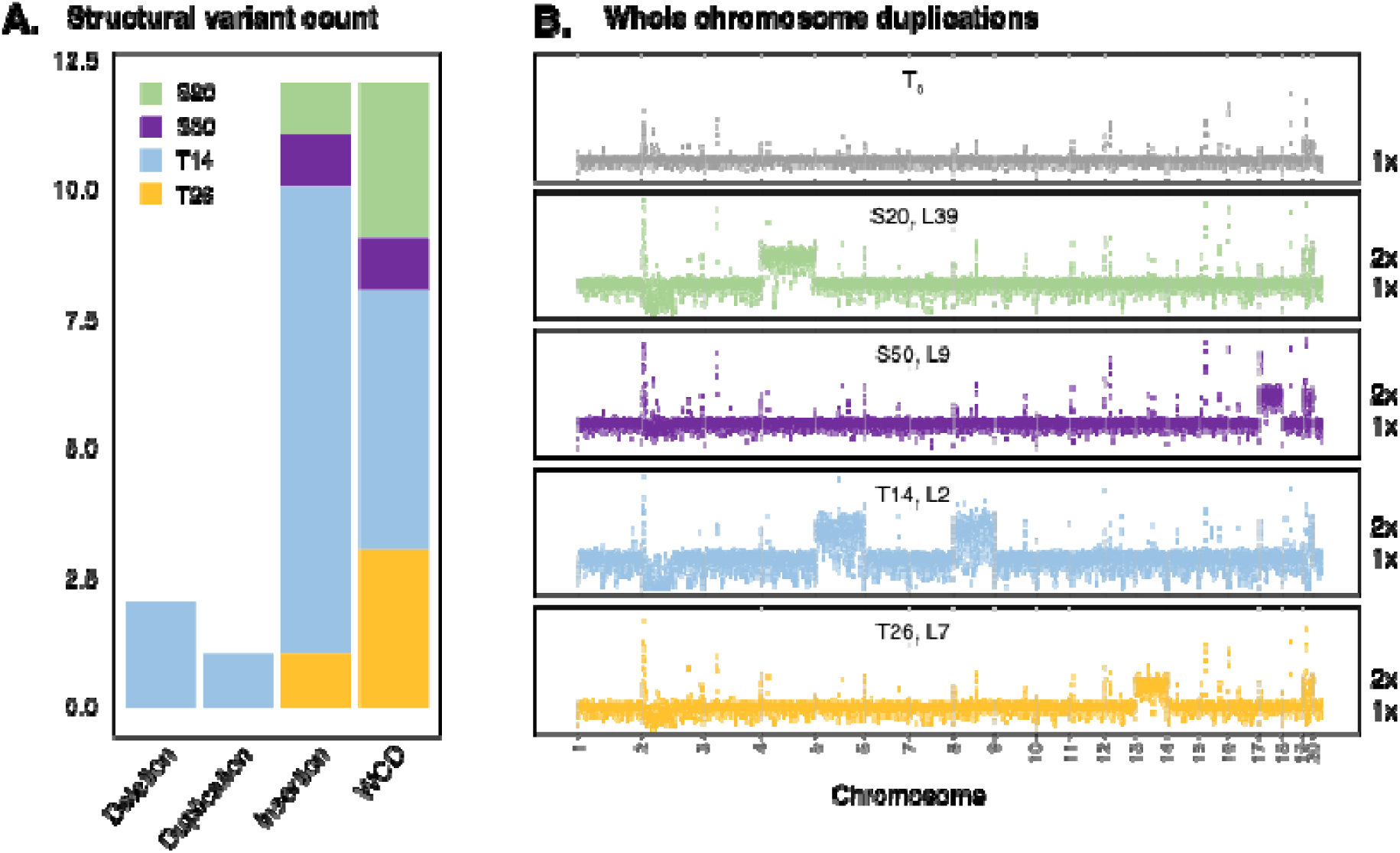
Structural de novo mutations (SM). **A.** distribution of the different types of structural mutations over the different MA conditions (Tables S8, S10). Although T14 has an *U_SM_* 4-7 times higher than all other MA, this only proves significant when compared to S50 (One-sided Fisher exact test, p-value = 0.014, Table S11). WCD: Whole chromosome duplication. **B**. Examples of WCD, as detected with raw Illumina sequencing depth (y-axis) per 1000 bp window of the nuclear genome in the ancestral line (*T_0_*). Chromosome numbers are displayed on the x-axis. WCD were observed in each MA condition (Fig. S3, Table S8).

### Structural mutations

Twenty-nine MA lines plus the *T_0_* were sequenced using long-read technologies (Table S9) to detect structural mutations (SM) with a combination of de novo assemblies and read- and assembly-based structural variant callers. A total of fourteen SM (two deletions, twelve insertions) were found, from 363 to 7214bp length. Eleven insertions and one deletion seemed to come from the activity of transposable elements (TE), with five different potential TE families identified (Table S10), containing transposon-like domains with transposase and endonuclease annotations.

Two other potential insertions were found in T26 line 26 and T14 line 16, respectively (Table S10); however, we were unable to confirm them by PCR because of multiple amplifications. Finally, using raw Illumina coverage, a duplication of the last quarter of Chr16 (±110kb) was detected in T14_39 (Fig. S4). Thus, a total of 15 different SM candidates were identified. Looking at the rate of SM per genome per generation (*U_SM_*) in each MA condition, T14 has an *U_SM_* 4-7 times higher than all other MA, although this only proved significant when compared to S50 (One-sided Fisher exact test, p-value = 0.014, Table S11).

### Fitness and *µ*

The evolution of fitness during MA experiments was evaluated using GR of MA lines as a proxy. Looking at the whole of the experiments, only in T26 the GR changed significantly over the course of the experiment (Spearman’s correlation, p-value = 0.014, rho = -0.63). However, in each experiment there were individual MA lines with a significant change in GR (Table S12). All such lines from T14 and T26 had a declining GR while all significant lines from S20 and S50 showed an increase in GR. In the lines with changing GR, the most common mutations were missense mutations (26/54) followed by intergenic mutations (14/54) and synonymous mutations (12/54); there was also 1 frameshift and 1 lost start mutation. Several lines also had structural mutation (SM) or whole chromosome duplications (WCD) but not all lines with SM or WCD had a change in GR (Tables S8, S10, S12). Although most lines with a significant change in GR had one or multiple mutations, this was not the case for T14_10. Thus, there is no evident correlation between fitness and mutation rate or type.

### DNA methylation in different conditions

To investigate whether epigenetic modifications between environments can explain a part of the variation in µSNM, DNA methylation level was estimated in 18 MA lines (Table S9, PromethION samples) for four different contexts (5mC, 5mCG, 4mC and 6mA, Table S13).

First, global genome methylation for each of the contexts were very similar between conditions (max. 0.5% variation between conditions), although statistically different (Chi-squared test, p-value 2.2×10^-16^): 23% for 5mCG, 12% for 5mC, 0% for 4mC and 0.5% for 6mA.

Second, there was no significant correlation between mutation and DNA methylation in any context in any condition (One-sided Fisher exact test, p-value > 0.06 for all) except for 5mC in S20 (One-sided Fisher exact test, p-value = 0.038), where methylated C had higher chance to mutate. However, mutations being very rare events, our statistical power is limited; the minimum difference in mutation rate between 5mC and unmethylated C we could have detected with our data is 2.4x for S20, 2.0x for S50, 2.4x for T14 and 1.9x for T26. There were no mutations on the few sites that carried 4mC and 6mA marks.

Third, there were mutations on methylated positions for 5mCG and 5mC marks in each MA, making it possible to compare methylation levels of mutated sites between MA experiments (Table S13). For 5mCG, methylation levels of mutated sites in T14 were significantly lower than in S20 and T26 (One-sided Fisher exact test, p-value < 0.05 for both); for 5mC, methylation levels of mutated sites in T14 were significantly lower than all others (One-sided Fisher exact test, p-value < 0.03 for all).

### The influence of transcription on mutation rate

Transcription may be mutagenic through transcription associated mutagenesis^27^ or protective through transcription coupled repair^28^. Thus, differences in transcription rate and landscape could in part explain difference in *µ* between conditions. We tested if mutated genes are differently expressed in MA conditions than non-mutated genes and found a significant difference in T26, where mutated genes are higher expressed (Table S14); however, the difference in median expression was quite small (4.16 vs 4.7, expression values in log2(RPKM+1)).

### Reactive oxygen species (ROS) in MA conditions

Oxidative stress due to ROS can cause damages to the DNA, particularly C→T^29–31^ and G→T^32^ mutations. To evaluate if this can explain part of the plasticity of *µ*, we measured general intracellular ROS using the H_2_DCFDA probe (Table S15, Fig. S5). Fluorescence geometric mean and median, calculated as the difference between each stained sample and a paired negative control, differed significantly among environmental treatments (Kruskal-Wallis p-values 0.009843 and 0.02096, respectively). S50 had the highest metric values across replicates (average geometric mean 0.00126, average median 0.00068; next highest values are in the Control condition with 0.000557 and 0.000402, respectively), whereas temperature and low-salinity treatments produced weaker and less consistent responses.

## DISCUSSION

### Explanations for the observed mutation rate and spectrum plasticity

A variety of explanations are possible for the plasticity of *µ*. First, a change in environment may cause stress-induced mutagenesis (SIM), which is well- documented in bacteria^33,34^, yeast^35^ and human cancers cells^36^. SIM directly increases *µ* through the expression of mutagenic DNA repair genes^33^ or low fidelity DNA polymerases^35,37^. SIM may be an explanation for the increase in *µ* in conditions were growth rate was significantly low (S50), indicating a possible stress for the cells. However, *O. tauri* did not show a growth reduction across all conditions of the MA experiments despite variations in *µ*.

Second, DNA methylation on C nucleotides has been shown to be mutagenic in bacteria^38^, animals^39–41^ and plants^42,43^. Under the effect of temperature and salinity variations, DNA methylation pattern and landscape may change^18,19,44–46^; thus, this might partially explain differences in mutation rate and spectrum between conditions. However, in *O. tauri* we see no clear correlation between *µ* and C methylation (Table S13). This is in line with earlier findings that showed that DNA methylation might not increase local *µ* in unicellular algae^47,48^.

Third, DNA transcription may impact *µ*. Assuming that transcription landscape and rate vary between environments, such variations may play on *µ* as found here in T26, although this was not detected in other conditions or previous studies^26^.

Fourth, reactive oxygen species (ROS)^49^ can cause oxidative damage to the DNA. Importantly, it creates 8-oxoG, which can be paired with A, causing G→T mutations, and C-deamination, which causes C→T mutations^29–31^. In S50, we observe higher ROS activity, which can, in part, explain the increase in *µ* and mutation bias.

Fifth, the accuracy of DNA polymerases can be affected by temperature^50^ and stressors such as heat shock^51,52^. For example, commercially available DNA polymerases from *E. coli* have a substitution rate up to 28 times higher at 37°C compared to 10°C^53^.

Sixth, salinity stress might demand energy to produce osmolytes^54^. Osmolytes are metabolites, easily soluble, neutral and non-toxic organic molecules that accumulate in the cell until osmotic equilibrium. Higher osmolyte production may limit resources for other functions linked to replication fidelity, DNA repair or ROS regulation.

Last, a noteworthy observation is the activation of TE in our MA lines in T14 (Fig.3, Table S10) indicating potential temperature responsive elements. The original discoverer of TE already suggested that they would be activated by stress^55^. In plants, many cases have been reported where transposons show activation after salt stress^56,57^, heat^58,59^ and particularly cold temperatures^60–62,62,63^.

### Consequences of a plastic *µ*

The fact that *µ* and spectrum change with the environment has several implications. First, many equations in population genetics depend on *µ* as a quantifiable parameter, but the mutation rate estimated in lab conditions may be far from the mutation rate in natural habitats. Since the natural environment is assumed to always be slightly stressful due to variations in resource availability, competition, predation, or abiotic factors, the mutation rate in the wild may be higher than that in the lab in most cases.

Second, efforts to explain variation in µ between strains or species should integrate the impact of the environment, or disentangle µ variations linked to external parameters.

Third, the mutation rate and spectrum define the amount and type of new genetic diversity available for selection; if they change with the environment, this has important consequences for evolutionary trajectories of different populations^64^. Thus, by playing on mutation rate and spectrum, environmental changes like global warming may directly impact the capacity of a species to adapt to these changes.

Finally, the mutation rate seems to be higher in individuals of poor fitness^65^ and in the majority of tested environmental changes (Table S1). A higher *µ* causes an increase of mutation load and thus a fitness decrease again, which, in turn, will further increase *µ,* and so on^66^.

### Mutation bias and GC-content evolution

The substantial variation in mutation bias may affect i) the distribution of fitness effects of mutations, ii) the genomic GC-content, and iii) *µ* itself. First, genomic GC-content is strongly influenced by mutation bias^67^, particularly at neutral sites. Populations exposed to markedly and long-term different ecological contexts may therefore experience divergence in the evolution of GC content. Second, codons ending in G or C are often favored over codons ending in A or T because of slight differences in selective coefficients^68,69^, a phenomenon known as codon usage bias^70^. A change in the mutation spectrum may therefore alter the proportion of mutations that convert a codon toward a new synonymous codon with either higher or lower selection coefficient. Third, because GC nucleotides mutate faster than AT nucleotides, genomes with higher GC content tend to exhibit higher mutation rates^26^.

In this way, a shift in GC content driven by a strong mutation bias may ultimately increase or decrease the average per-nucleotide mutation rate of a given genome.

## MATERIALS & METHODS

### Mutation accumulation experiment

Four different mutation accumulation (MA) experiments were performed on *Ostreococcus tauri* strain RCC4221 using a flow cytometry protocol described previously for liquid medium^71^. The identity of the strain was confirmed by 18S rDNA PCR coupled with Sanger sequencing at the start of the experiment. Cells were cultivated in L1 medium generated from sterile natural sea water (Bigelow L1 Culture Medium Kit) in a light:dark cycle of 12:12 hours. The ancestral population used for the MA experiments (*T_0_*) was started with an estimated single cell isolated by dilution after measuring cell density by flow cytometry. Then, the *T_0_* was separated in 4 flasks and left 3 weeks for acclimation to the MA conditions: low salinity: S=20g/L (‘S20’) and high salinity: S=50g/L (‘S50’), both at 20°C; and low temperature: T=14°C (‘T14’) and high temperature: T=26°C (‘T26’), both with a salinity of 33g/L. The changes in salinity were effectuated by diluting natural sea water at 33g/L with milli-Q water or by adding artificial sea water salts (for 1L, add: 0.855g NaCl, 0.26g KCl, 1.42g MgCl_2_.6H_2_O, 0.51g CaCl_2_.2H_2_O, 2.1g MgSO_4_.7H_2_O, 0.07g NaHCO_3_).

The cell density of each *T_0_* culture was measured again by flow cytometry to sample the volume corresponding to 50 cells, which were diluted in 50 mL culture medium and then used to inoculate one 48-well plate with 1 mL (∼1 cell per well); this was repeated for all plates (six 48-well plates per MA). In addition to MA lines, a control culture with 100 cells/well was initiated in 12 replicates for each MA condition. From this initial single cell inoculation, MA lines were inoculated in 48-well plates in 6 replicates each (1 row per MA line); the 6 replicates limit the loss of MA lines due to technical errors or lethal mutations^71^. Single cell bottlenecks were then performed every 14 days to reduce selection to a minimal level. At each bottleneck time, cell density of MA line populations was measured by flow cytometry and the first well with living cells found was used for reinoculation with statistically 1 cell per well, by adding the volume corresponding to 10 cells to 10 mL of medium and using that to re-inoculate 6 wells. The control culture was reduced to 100 cells/well every bottleneck using the same method. The bottlenecks were repeated 24 times for S20 and S50, 19 times for T14 and 15 times for T26. In T26 there were some MA line losses; thus, new MA lines were inoculated from existing MA lines. These so called ‘sister lines’ thus share a number of generations and, potentially, mutations (Table S16). The cell density was also used to estimate the average daily growth rate *GR* between each bottleneck using the following formula:

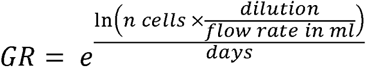

Thus, the number of generations between each bottleneck equals *Gen* = (*GR*-1) x 14. The average *N_e_* per MA line between each bottleneck was calculated using the harmonic mean of cell number each day, starting from 1 cell to the final cell number, with the number of cells per day calculated with *GR*. The *N_e_*in each well never exceeded 8 cells. The *GR* was also used as a proxy of MA line fitness during the experiment. For fitness tests, the *GR* was normalized by dividing the *GR_MA line_* by the *GR_control_*for each bottleneck.

At the end of the MA experiment, MA line populations were transferred to 30 mL flasks and allowed to grow for 2 to 3 weeks in MA conditions in order to obtain sufficient cell numbers for DNA extraction. The final times of MA S50 were grown in 33g/L medium, to boost growth. High molecular weight DNA was extracted from 155 MA lines plus the ancestral line *T_0_* using a CTAB extraction protocol modified from^72^. Total DNA libraries were prepared and sequenced by the Joint Genome Institute (JGI, Berkeley Lab, California) with Illumina NovaSeq 4000 in 150 paired-end reads.

### Nucleotide and short indels de novo mutation identification

Raw Illumina reads were trimmed at the JGI and the read quality was checked using FastQC v0.12.1^73^. Reads were mapped against the reference genome (GCF_000214015.3^74^) with bwa-mem2 v2-2.1^75^ and the resulting BAM files were treated using samtools v1.19^76^. The callable genome (*G\**) per MA was defined through 3 steps: (i) samtools depth was used to keep the positions with at least 10 coverage in MA lines and 20 in *T_0_*, considering both base and mapping quality thresholds of 20 (options -q 20 -q 20); (ii) positions with a coverage of more than 2x the average depth were removed to exclude erroneous mappings in highly repetitive regions; (iii) only the positions that fulfilled these criteria in all samples of the same MA plus the *T_0_* were kept as callable sites. Variant calling was done with HaplotypeCaller from GATK v4.4.0.0^77^ with the *-ploidy 1* option. The resulting VCF files were compiled using samtools mpileup v1.18. Next, we filtered candidate de novo mutations by the following criteria: (i) the site is in *G\**; (ii) no reads with the same candidate in the ancestral line *T_0_*; (iii) maximum 3 reads with the same candidate in maximum 3 other MA lines (with an exception for sister lines, Table S16). All insertion-deletion (indels) candidates were manually reviewed using IGV v.2.15.1^78^. Primers were designed with Primer3^79^ to check all candidate mutations with 2 or more similar reads in other MA lines by Sanger resequencing, as well as all indels. For the SNM, 41 of 52 chosen positions were successfully resequenced, for indels this was 13 out of 19 (Table S17 for primers). Sanger sequencing showed a true positive rate of 100%. Thus, our criteria for candidate positions were strict enough and all retained candidate mutations were considered true. Then, SnpEff v.5.1^80^ was used to identify mutation effects. The total mutation rate as well as mutation rates per type of mutation were calculated using the following formula:

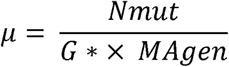

Where N_mut_ is the number of mutations, G* is the number of callable sites, and MA_gen_ is the sum of generations of all MA lines in a same experiment.

For the analysis of mutation spectrum, mutation rates for each of the 6 possible SNM were estimated (Fig. S1). We also looked at changes in genome composition using *µ_GC_*_→_*AT* and *µ_AT_*_→_*GC* (Table 1), which are the number of GC to AT mutations per generation per number of GC bases in the callable genome and the number of AT to GC mutations per generation per number of AT bases in the callable genome, respectively. With these rates we can calculate the equilibrium GC-content at which nucleotide composition is stable if we consider mutational process alone, or, at which the number of mutations from GC to AT and AT to GC are equal: *GCeq=µAT*→*GC/(µGC*→*AT µAT*→*GC)*.

### Detection of structural de novo mutations

Resequencing with long read sequencing was performed on 32 MA lines plus the T0 culture, using either PacBio HiFi or Nanopore PromethION, or a combination of PacBio HiFi and Nanopore MinION (Table S9); for the T0 and the 6 MA lines in this last category, reads were merged before assembly.

All samples with nanopore reads were adapter-trimmed using Porechop v0.2.4^81^ and NanoFilt v2.8.0^82^, and read quality was assessed with NanoPlot v1.44.1^82^. Then, for all strains, three independent de novo assemblies were generated using Canu v2.3^83^, Flye v2.9.6^84^, and Hifiasm v0.24.0^85^. Each assembly was then polished separately with Racon v1.5.0^86^ and Medaka v2.1.0^87^, yielding six polished assemblies per strain. These long-read–polished assemblies were further polished with Pilon v1.24^88^, using the corresponding paired-end Illumina short reads. Assembly quality was evaluated using QUAST v5.3.0^89^, and large-scale collinearity with the reference genome was inspected using MUMmer4 v4.0.1 dotplots^90^. The final assembly retained for downstream analyses was selected based on contiguity, completeness, and chromosomal concordance with the newest reference genome^91^ (Table S9).

Before structural variant (SV) discovery, retained assemblies were filtered to keep only contigs aligning to the reference genome with ≥80% sequence identity and ≥30 kb cumulative aligned length, using MUMmer4 nucmer (-l 100 -c 500). SVs were then identified using six complementary approaches. Two were assembly-based: (i) minimap2 v2.28^92^ alignment of the filtered assembly against the reference followed by SVIM-asm v1.0.3 in haploid mode^93^, and (ii) nucmer alignment followed by MUM&Co^94^. Four were read-based: (iii) minimap2 alignment of Nanopore reads against the reference followed by SVIM v2.0.0^95^, (iv) DeBreak v1.3^96^, (v) SVDSS v2.1.0 (smooth, search, and call functions)^97^, and (vi) Sniffles2 v2.7.2^98^. To preserve SV detection in repetitive regions while also retaining confidently unique mappings, we kept reads with either mapping quality 0 (typically repetitive or multi-mapping reads) or ≥30 (high-confidence unique alignments), while excluding intermediate mapping qualities. Across callers, we required a minimum SV size of 50bp and at least five supporting reads.

To obtain a high-confidence consensus SV set, only variants detected by at least two independent callers were retained. For Nanopore sequenced and mixed strains, SV calls were merged across methods when breakpoints occurred within 30bp and SV lengths agreed within either 2% relative difference or 50bp absolute difference; for HiFi samples, calls were merged if they occurred within 20bp and SV lengths agreed within either 5% relative difference or 50bp absolute difference. All final SV candidates were tested by PCR using primers designed with Primer3^79^ around the mutated position (Table S17), allowing for direct verification of the SV by amplicon size on agar gel migration.

Finally, we looked at the raw genomic coverage with bedtools v2-2.18.0^99^, using a window size of 1kb. This analysis provides evidence of large duplications in several MA lines, of which the majority are whole chromosome duplications (WCD) (Fig. 3, S3, Table S8); 2 MA lines with WCD were also sequenced by long read technologies and duplications were confirmed with long read raw coverage.

### Methylation analysis

To see if differential DNA methylation contributes to the differences in mutation rate between conditions, we looked if there was a correlation between local *µ_SNM_*and 4 different DNA methylation marks: 5mCG, 5mC, 4mC and 6mA. To this end, 3 to 5 MA lines from each condition for a total of 18 samples were long read sequenced by Nanopore Technology (Table S9). For S50, 3 intermediary times (‘IT’) were sequenced instead of the final time of the MA experiment, because the final cultures were amplified in a flask with normal medium instead of high salinity medium, which might have changed methylation status. Modified base-calling and alignment of long Nanopore reads were done with Dorado v.0.9.1^100^, using the dna_r10.4.1_e8.2_400bps_sup@v5.0.0 model with 4 different DNA methylation models (5mCG_5hmCG, 5mC_5hmC, 4mC_5mC and 6mA, all v3). BAM files were sorted and indexed with samtools v1.19 and converted to bedMethyl files using Modkit v0.3.1^101^. The default thresholds set by Modkit were quite low (0.703-0.718 for 5mCG, 0.574-0.598 for 5mC, 0.508-0.516 for 4mC, 0.783-0.791 for 6mA) but maintained, since increasing the *--filter-threshold* can introduce a bias, artificially increasing methylation levels as base probability is generally higher for methylated than for canonical bases. We analyzed methylation levels in all samples combined per condition to get a more confident methylation call, using only the positions covered ≥5x times^102^. Positions were considered methylated if ≥80% of reads supported the modification in the majority of samples. To evaluate the minimum difference in mutation rates that could be detected with our data between methylated and unmethylated C, we used the standard error of the log-transformed rate ratio (RR), which was estimated with 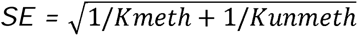, where *Kmeth* and *Kunmeth* are the observed numbers of mutations at methylated and unmethylated C, respectively. This allowed to calculate the RR values for which |ln(RR)| > 1.96 x SE, corresponding to a detectable difference at α = 0.05.

### Transcription analysis

RNA was extracted from *O. tauri* cultures grown in the 4 experimental conditions plus standard lab conditions (T=20°C, S=33g/L) as a control, for both day and night transcriptomes in 3 technical replicates for a total of 30 samples, although 2 failed sequencing (1 replicate of S50 night and 1 of T14 night). Extraction was done with the Monarch Total RNA Miniprep Kit. Total RNA libraries were prepared and sequenced by the Joint Genome Institute (JGI) at Berkeley Lab, California using a NovaSeq S4 with 150bp paired-end reads. Raw reads were filtered at the JGI, deinterleaved using Seqtk v1.3 and mapped against the reference genome using STAR v2.7.10b with the options ‘–alignIntronMin 20’ and ‘—alignIntronMax 500000’. For the correlation between mutability and gene transcription, gene expression levels were quantified using the RPKM method in R v4.4.3 with the help of the edgeR package v4.4.2^103^.

### H_2_DCFDA assay for ROS activity

To see global intracellular ROS activity, *O. tauri* cells were incubated with 40µM H_2_DCFDA in MA conditions in the dark for 45 minutes. This probe diffuses through the cell membrane and becomes trapped after de-acetation by intracellular esterases, upon which it can interact with ROS and become fluorescent^104^. After incubation, cells were washed once with, and resuspended in, salinity adjusted 0.2µm filtered sea water, immediately followed by cytometric measures of green fluorescence. For each replicate in each condition a paired negative control, incubated without adding H_2_DCFDA, was included. This protocol was repeated 6 times on the same day, from 10am to 4pm. ROS values (geometric mean, median) were calculated as the difference between the sample and its paired negative control.

## Supporting information

All supplemental tables

All supplemental figures

## ACKNOWLEDGEMENTS

We acknowledge the GenoToul Bioinformatics platform (Toulouse, France) for bioinformatics analysis support and cluster availability and all members and former members of the GENOPHY team. We also thank the BIOPIC platform for support with the flow cytometry, the BSBII platform and especially Vladimir Daric for help with data analysis, the Bio2Mar platform and especially Nyree West for support with experimental procedures and the glassware cleaning staff for their great help in avoiding contaminations. The work (proposal: 10.46936/10.25585/60008631) conducted by the U.S. Department of Energy Joint Genome Institute (https://ror.org/04xm1d337), a DOE Office of Science User Facility, is supported by the Office of Science of the U.S. Department of Energy operated under Contract No. DE-AC02-05CH11231. This work was funded by the ANR ANR-23-CE20-0013 and ANR-21-CE02-0026.

## AUTHOR CONTRIBUTIONS

LM performed the mutation accumulation experiments, analyzed the data and drafted the first version of the manuscript. ALi, CV and CE performed initial bioinformatic analyses on the raw reads. FS and ALa contributed to experimental procedures. KB and IG managed sequencing. GP and MK managed the project, MK contributed to the mutation accumulation experiment and writing. All authors contributed to the final version of the manuscript.

## SUPPLEMENTARY MATERIAL

**Table S1 *µ* plasticity in the literature**

**Table S2 Additional *µ* information**

**Table S3 Details of MA lines**

**Table S4 Callable genome (*G\**) details per chromosome and per MA**

**Table S5 Details of de novo mutations**

**Table S6 Mutations and callable genome size (*G\**) for compartments of the nuclear genome per MA**

**Table S7 Significance of differences in *µ_SNM_***

**Table S8 Whole chromosome duplication (WCD) rate**

**Table S9 Long read sequencing**

**Table S10 Structural mutations (SM)**

**Table S11 Comparison of SM rates**

**Table S12 MA lines with a significant change in growth rate over the course of the experiment**

**Table S13 DNA methylation**

**Table S14 Transcription and mutation rate**

**Table S15 ROS assay for oxygenic stress**

**Table S16 Sister lines in T26**

**Table S17 Primers used to check de novo mutations**

**Fig. S1 Details of the SNM spectrum**

**Fig. S2 The correlation between *µ_C:G_***_→_***T:A* and total *µ_SNM_***

**Fig. S3 All whole chromosome duplications (WCD)**

**Fig. S4 Sequencing depth of the last half of Chromosome 16 in T14 L39, showing a large duplication**

**Fig. S5 Effect of MA conditions on ROS production in *O. tauri***

**Fig. S6 Principal component analysis (PCA) of gene expression in *O. tauri* under different conditions – dimensions 1 and 2**

**Fig. S7 Principal component analysis (PCA) of gene expression in *O. tauri* under different conditions – dimensions 2 and 3**

