## Supplementary material for "Plasticity of the mutation rate and spectrum in eukaryotic marine phytoplankton": All supplemental figures

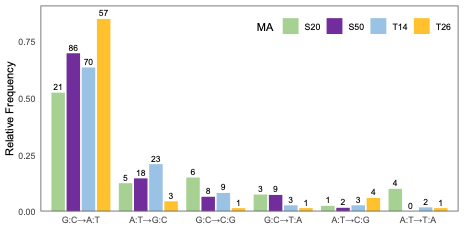
**Figure S1.** SNM spectrum

Transitions (G:C→A:T and A:T→G:C) were the most common SNM in any condition and this bias gets stronger with increasing *µ_SNM_*. In high temperature (T26) 83% of all SNM are G:C→A:T transitions.

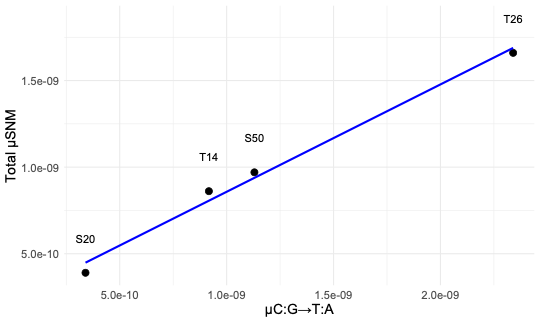

**Figure S2.** The correlation between *µ_C:G_*_→_*_T:A_* and total *µ_SNM_*

With increasing *µ_SNM_* in the different conditions, *µ_C:G_*_→_*_T:A_* is the only type of SNM that shows a significantly increased rate. The correlation between *µ_C:G_*_→_*_T:A_* and *µ_SNM_* is linear (adjusted R² = 0.985, p = 0.005).

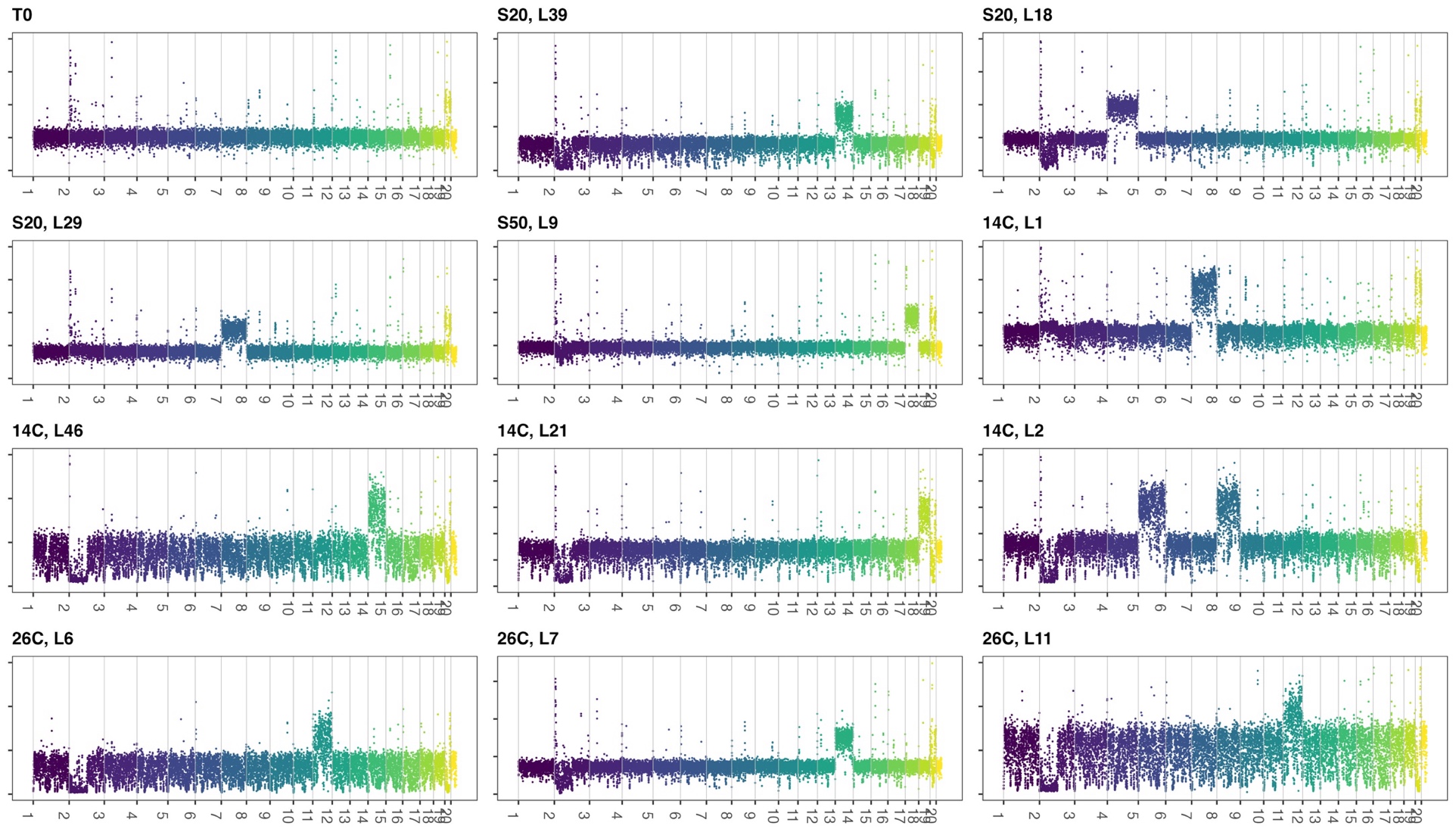

**Figure S3. All whole chromosome duplications (WCD)**

Illumina sequencing depth (y-axis) per 1000 bp window of the nuclear genome in the ancestral line (*T_0_*) and MA lines with one or multiple WCD. Chromosome numbers are displayed on the x-axis.

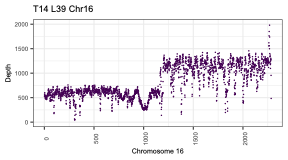

**Fig. S4** Sequencing depth of the last half of Chromosome 16 in T14 L39, showing a large duplication

Raw Illumina sequencing depth is plotted from position 299900 to the end of chromosome 16 (528281), showing a duplication of almost 110kb, starting at position 419300. It is uncertain if and where the duplicated sequences integrated since only short-read data are available for this line.

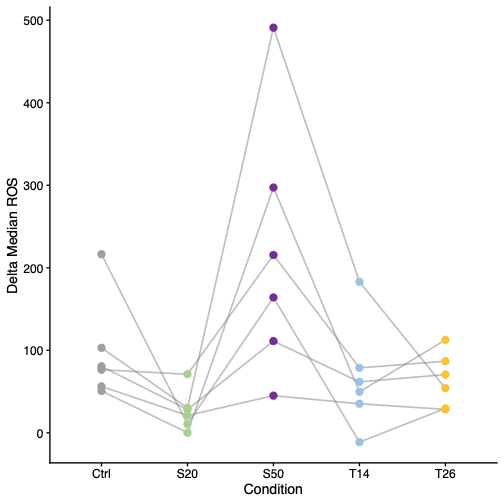

**Fig. S5** Effect of MA conditions on ROS production in *Ostreococcus tauri*

The H2DCFDA probe was used to measure intracellular stress in Ostreococcus tauri cells in all 4 MA conditions, plus standard lab conditions ("Control", 20 C, 33g/L salt) (Table S15). Using a flux cytometer, the Geometric Mean of green fluorescence in a population of 5000-5600 cells for each point was measured. For each point, a negative population was measured as well, which had not been incubated with the probe but followed an otherwise identical protocol. This allowed to take out background signal by calculating the Δ Geometric Mean. The measures were repeated six times over the course of a single day, showing a large variation with time.

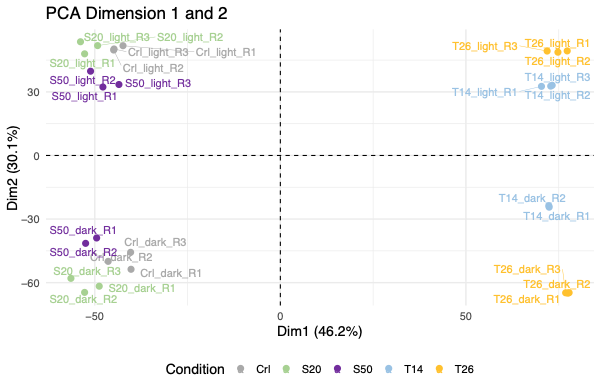

**Fig. S6** Principal component analysis (PCA) of gene expression in *O. tauri* under different conditions – dimensions 1 and 2

RNA was extracted from cultures in each of the 4 MA conditions plus standard laboratory conditions (temperature = 20°C, salinity = 33g/L), for each condition both day and night transcriptomes, all in triplicate. However, one T14 night sample and one S50 night sample failed sequencing. Gene expression was normalized using the trimmed mean of M values (TMM) and the data were visually examined by plotting the PCA, where the first axis explains 46.2% and separates the control/S20/S50 from T14/T26, the second axis explains 30.1% and separates day from night samples.

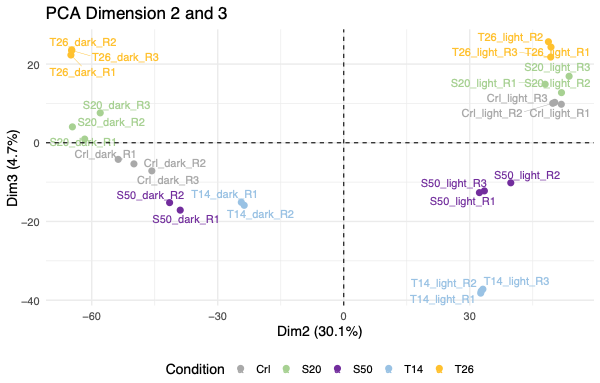

**Fig. S7** Principal component analysis (PCA) of gene expression in *O. tauri* under different conditions – dimensions 2 and 3

See the legend of Fig. S5. In the PCA, the second axis explains 30.1% and separates day from night samples, and the third axis explains 4.7% and roughly separates T26/S20 from T14/S50.
